# Nucleosome dynamics of human iPSC during the early stages of neurodevelopment

**DOI:** 10.1101/398792

**Authors:** Janet C Harwood, Nicholas A Kent, Nicholas D Allen, Adrian J Harwood

## Abstract

Regulation of nucleosome positioning is important for neurodevelopment, and mutation of genes mediating chromatin remodelling are strongly associated with human neurodevelopmental disorders. Unicellular organisms possess arrays of highly positioned nucleosomes within their chromatin, occupying up to 80% of their genomes. These span gene-coding and regulatory regions, and can be associated with local changes of gene transcription. In the much larger genome of human cells, the roles of nucleosome positioning are less clear, and this raises questions of how nucleosome dynamics interfaces with human neurodevelopment.

We have generated genome-wide nucleosome maps from an undifferentiated human induced pluripotent stem cell (hiPSC) line and after its differentiation to the neuronal progenitor cell (NPC) stage. We found that approximately 3% of nucleosomes are highly positioned in NPC. In contrast, there are 8-fold less positioned nucleosomes in pluripotent cells, with the majority arising *de novo* or relocating during cell differentiation. Positioned nucleosomes do not directly correlate with active chromatin or gene transcription, such as marking Transcriptional Start Sites (TSS). Unexpectedly, we find a small population of nucleosomes that remain positioned after differentiation, occupying similar positions in pluripotent and NPC cells. They flank the binding sites of the key gene regulators NRSF/REST and CTCF, but remain in place whether or not their regulatory complexes are present. Together, these results present an alternative view in human cells, where positioned nucleosomes are sparse and dynamic, but act to alter gene expression at a distance via structural conformation at sites of chromatin regulation, not local changes in gene organisation.

## Main

Nucleosome maps generated by MNase-seq show the dynamic nature of nucleosome positioning. Narrow sequence read mid-point distribution profiles are indicative of highly positioned nucleosomes present at the same position in all cells of the population, whereas broader distributions occur where there is substantial variation in nucleosome positioning between individual cells (Figure 1A). By mapping nucleosomes in the same human induced pluripotent stem cell (hiPSC) line in both its undifferentiated, pluripotent cell state and following differentiation to the neural progenitor cell (NPC) stage, we followed the changes in nucleosome positioning during early stages of neurodevelopment. We found that only 2.7% of nucleosomes are highly positioned in human NPC (408,152 of a theoretical total of 15 million based on an assumption of 1 nucleosome per 200 bp). This is consistent with observations from other human and mammalian cells [1,2], but contrasts with the yeasts and *Dictyostelium* genomes where positioned nucleosomes occupy approximately 80% of the genome [3–6]. Using the same analysis on data generated from the human lymphoblastoid cell lines, K562 and GM12878 [7] we calculated that 2.4% and 1.6% of nucleosomes respectively were highly positioned in these cell lines (Supp Table S1).

**Figure 1:**
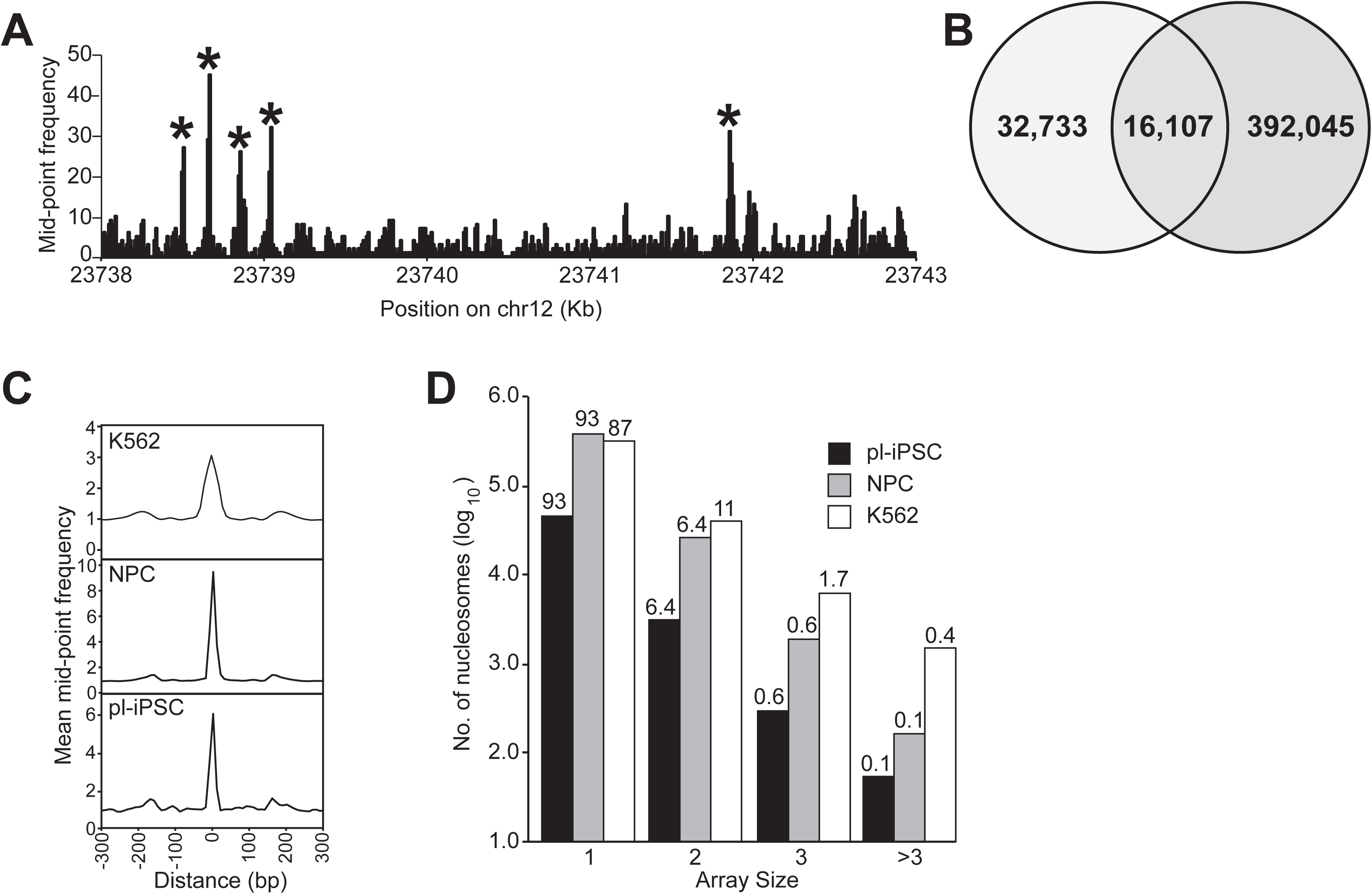
Nucleosome dynamics of human pluripotent stem cell and differentiated cells. A: Genome-wide nucleosome maps were generated from the frequency distribution of the midpoint read positions of pair-end sequenced MNase-resistant fragments in the size range 138-161 bp (spanning the nucleosome footprint). Figure shows a section of a nucleosome map taken from chromosome 12 derived in pluripotent iPS cells to show examples of highly positioned nucleosomes (*). B: Venn diagram showing the overlap in the genomic location of highly positioned nucleosomes in pluripotent iPSC (pl-iPSC) and in the same cell line differentiated into neural progenitor cells (NPC). C: Average frequency distribution of nucleosome positions relative to each other. Data was aligned to each mapped nucleosome in the genome and plotted for a 600 bp window for pluripotent iPSC (pl-iPSC), iPSC-derived NPC (NPC) and the chronic myelogenous leukaemia (Cml) derived cell line, K562. In all cases, a prominent single nucleosome is presented with only minor flanking peaks, indicating that the majority of nucleosomes do not occur as long arrays in human cells. D: Distribution of nucleosome array sizes for pluripotent human pl-iPSC, NPC and K562, calculated as the number of nucleosomes within a distance of 150-200 bp of each other. Bars show the number of nucleosomes arrayed as singletons (1), pairs (2), triplets (3) and greater than 3 (>3), displayed as log_10_ values and with the percentage of the distribution within a cell line is shown above the column.

In contrast, there were 8.4-fold fewer positioned nucleosomes (0.33% of the total nucleosome number) in the pluripotent cell state, indicating that a substantial increase in positioned nucleosomes occurs during differentiation from pluripotent cells to NPC. Comparison of both maps shows that 32% of the positioned nucleosomes present in the pluripotent hiPSC, retain their position during differentiation to NPC (Figure 1B). This indicates that there are considerable changes in nucleosome positioning during human neurodevelopment, but also that there are a small population of nucleosomes that remain positioned, albeit corresponding to only approximately 0.1% of the total nucleosome population.

To determine how human nucleosomes are distributed throughout the genome, the chromatin structure surrounding each positioned nucleosome was analysed in pluripotent hiPSC and NPC cell states, and for the K562 cell line. The chromatin structure had a broadly similar pattern, with the majority of nucleosomes present as singletons, and few nucleosomes (7.0%, 6.9% and 13% respectively) present as pairs or nucleosome arrays (Figure 1C & D). This indicates that the basic pattern of nucleosome positioning is similar across different human cell types.

We examined the distribution of positioned nucleosomes across different chromatin states. Using the 15 state model of Ernst and Kellis [8,9], which classifies chromatin from highly active to repressed states based on modifications, functions and associated proteins, we examined whether positioned nucleosomes partition into particular chromatin states. We found no significant enrichment for the major chromatin states, either transcriptionally active, repressed or heterochromatin, with the exception of chromatin state 8, which is characterised by its association with the chromatin architectural protein CCCTC-Binding factor (CTCF) [10] (Figure 2 A&B).

**Figure 2:**
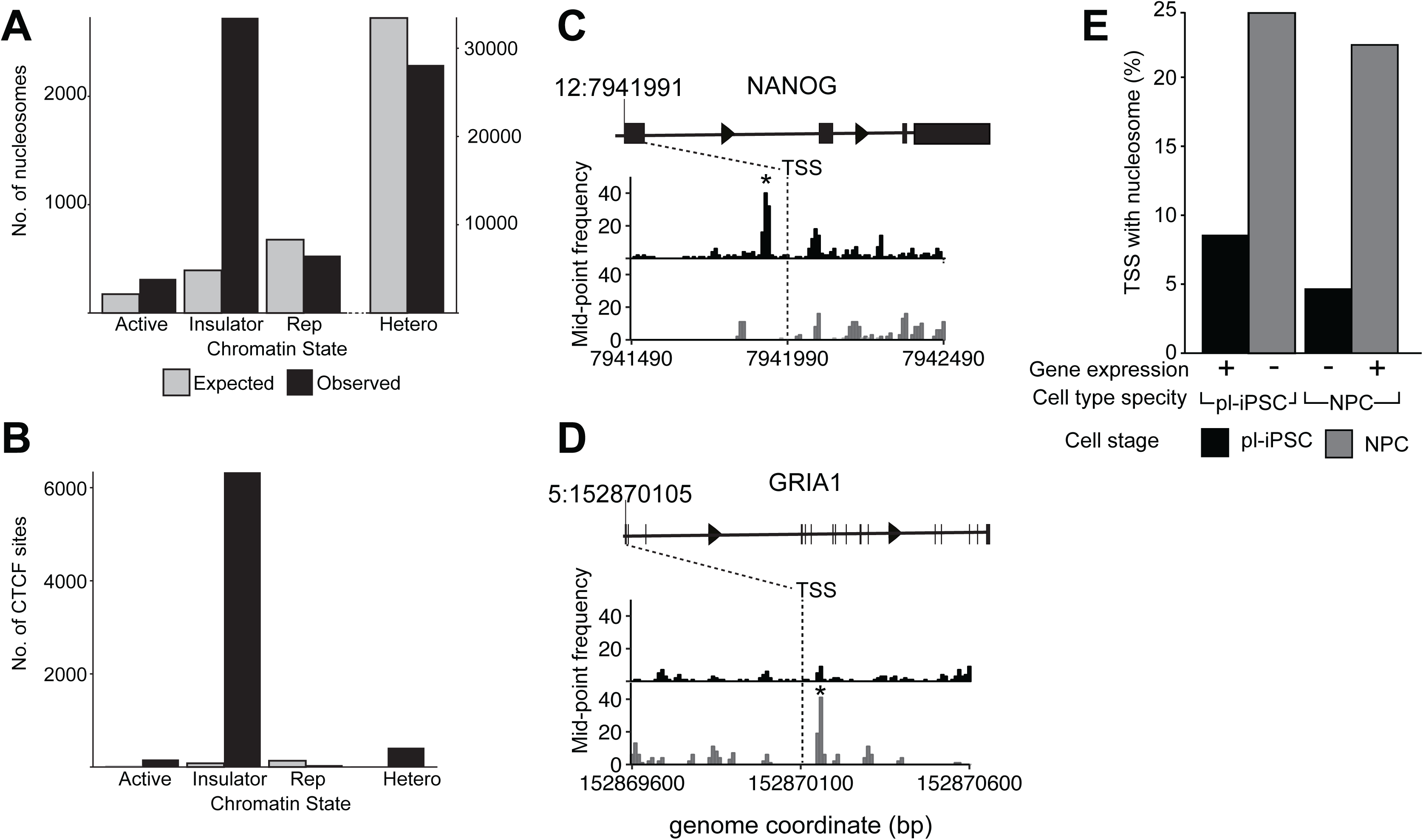
Highly positioned nucleosomes, chromatin states and transcriptional activity. A: Bar chart showing the number of nucleosomes that map to selected chromatin states [8]. Active (State 1); Insulator (State 8); Repressed, Rep, (State 12) and heterochromatin, Hetero (State 13). Shown are the observed number of nucleosomes per chromatin state (black) and the expected number calculated assuming a random distribution across the genome (grey). B: Bar chart showing the number of CTCF sites that map to different chromatin states, using the same methodology as A. C, D: Nucleosome maps at and surrounding, the TSS region of the (C) NANOG and (D) GRIA1 genes in pl-iPSC (upper, black) and in NPC cells (lower, grey). The TSS is indicated by a dashed line at chr 12: 7,941,991 for NANOG and chr5: 152,870,105 for GRIA1. * indicates the position of a highly positioned nucleosome associated with the TSS of the active gene. E: Genome-wide association of nucleosomes at TSS with gene activity. Based on RNA-seq data, TSS were identified that are associated with unique gene expression restricted to either pl-iPSC (n=3,833) or NPC (n=2,082). The number of TSS where a positioned nucleosome is present within ± 300 bp of the site (shown as a percentage of total unique TSS number) was calculated for the active gene (pl-iPSC TSS in pl-iPSC and NPC TSS in NPC) and the inactive gene (pl-iPSC TSS in NPC and NPC TSS in pl-iPSC). If positioned +1 nucleosomes were a strong predictor of gene activity, high numbers of nucleosomes would be expected for pluripotent specific genes in pl-iPSC, but not NPC stages, and for NPC specific genes in NPC, but not pl-iPSC stages. No correlation of between gene activity and positioned nucleosomes was observed.

The lack of enrichment of highly positioned nucleosomes within transcriptionally active chromatin states was unexpected given the known association of well-organised nucleosome positioning within active gene loci of non-mammalian cells and the association of gene expression with SNF-2 family chromatin remodelers, which control the placement of nucleosomes on DNA [11]. Therefore we investigated the local distribution of nucleosome positioning at Transcriptional Start Sites (TSS). In non-mammalian cells, actively transcribed genes are observed to possess a positioned nucleosome at their TSS, termed the + 1 nucleosome. In the human genome, we identified individual loci where a highly positioned nucleosome was present at the +1 position in the pluripotent cell state but not in NPC, for example the pluripotent specific gene NANOG (Figure 2C). Conversely we found individual loci where a highly positioned nucleosome was present at the +1 position only in NPC, such as the glutamate receptor subunit GRIA1 (GluR-1) (Figure 2D). However, this correlation did not hold at the whole genome level.

By cross-comparison of published RNA-seq data from human cells [12] with the non-redundant list of 83,179 mapped human TSS, we created datasets corresponding to the TSS positions of human genes that are transcribed uniquely in pluripotent or NPC cells. These datasets were then searched for the presence or absence of a positioned nucleosome within ± 300 bp of the TSS. These data showed that for genes expressed only in pluripotent cells there were more nucleosomes positioned at the TSS of genes in the inactive, NPC state than on genes with active expression (Figure 2E). For genes expressed only in NPC cells there was again a substantial number of nucleosomes associated with the TSS of the inactive gene, in the pluripotent cell state (Figure 2E). Overall, positioned nucleosome numbers were substantially higher in NPC cells at both active and inactive gene TSS reflecting the 8-fold general increase of positioned nucleosomes seen in NPC. These observations suggest that unlike other nucleosome modifications [13,14], the presence of highly positioned nucleosomes at TSS is not a genome-wide predictor of gene activity in human cells. To widen our analysis further, we examined the patterns of nucleosome positions at selected transcription factor (TF) binding sites involved in neurodevelopment, YY1, ATF2, and PAX6 (Figure S1), [15–17]. Again we observed no consensus pattern of positioned nucleosomes associated with these TF binding sites.

In contrast, well-positioned nucleosomes were present at the RE1 binding site [18] of NRSF/REST (Figure 3). These sites were selected by mapping the consensus sequence of RE1 sites to NRSF/REST ChIP-seq data [19] creating a dataset of 871 well-characterised RE1 sites. Aligning nucleosome positioning data to these RE1 sites across the genome revealed a consensus pattern comprising an array of nucleosomes flanking the RE1 site. NRSF/REST is transcriptional repressor that acts at a distance by scaffolding a protein complex on DNA, which contains repressive chromatin modifiers, including the H3K9 dimethyltransferase G9a/EHMT2, the histone deacetylase complex Sin3/ HDAC1/2, and the H3K4 demethylase LSD1 as well as the chromatin remodeller Brg1/SMARCA4 [20]. This complex is present at RE1 sites in mouse pluripotent cells [21] but it is lost upon neuronal differentiation. It was therefore unexpected to find that the pattern of nucleosomes surrounding the RE1 site in human pluripotent cells unchanged in NPC (Figure 3A). To analyse this pattern in more detail, we stratified sequencing reads into the size range 122-137 bp. This subset contains protected fragments for transcription factors or regulatory complexes with large DNA footprints, and identified a distinct peak of protected fragments at the RE1 site in pluripotent cells that corresponds to the NRSF/REST complex. This peak is reduced by more than 90% in NPC cells (Figure 3B).

**Figure 3:**
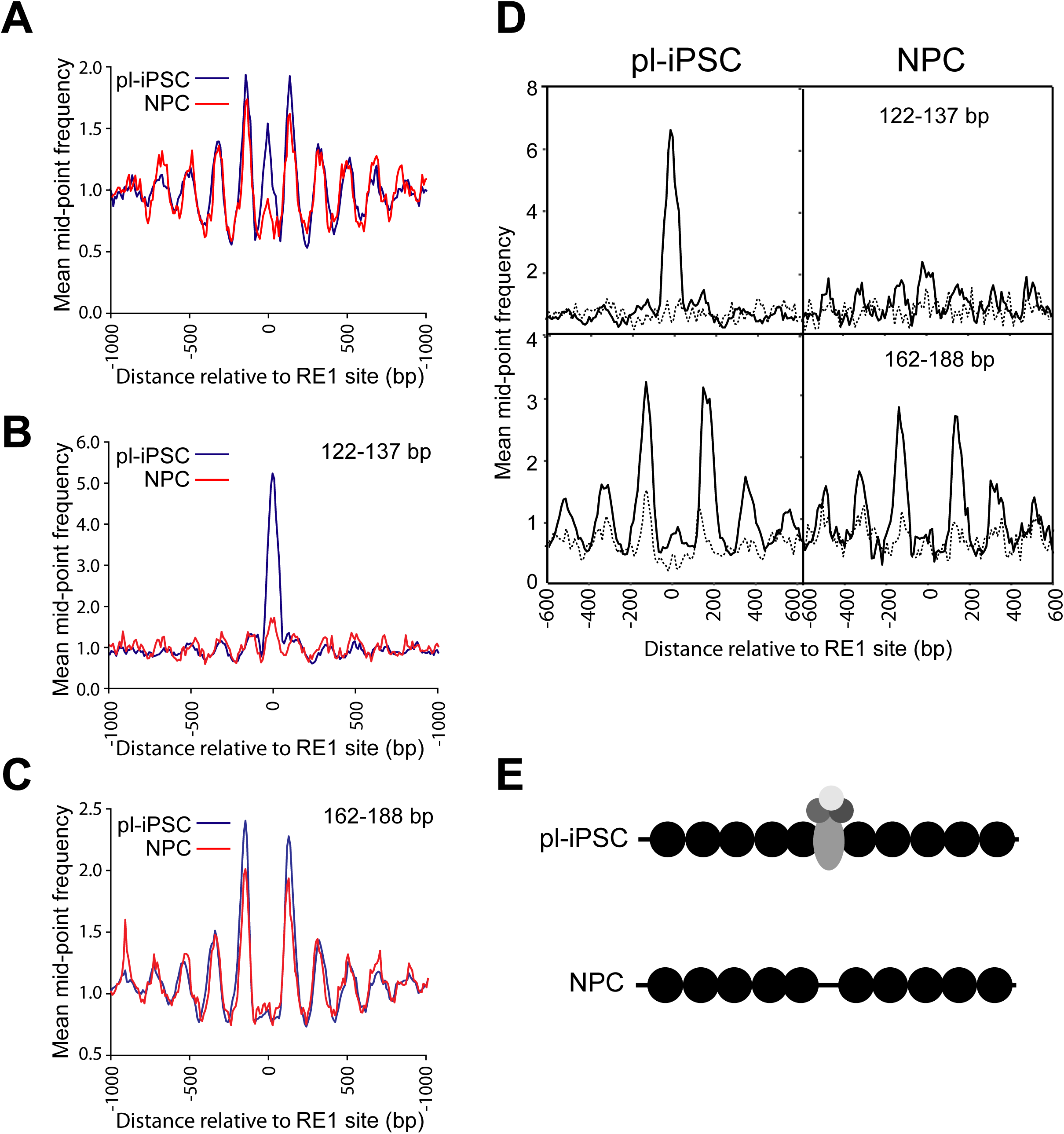
Nucleosome patterning associated with the RE1 NRSF/REST binding motif. A: The average distribution of nucleosomes centred on the RE1 site (n=871), spanning 2 Kb of flanking genomic sequence. B, C: Aligned pair-end sequence read data were stratified into two size classes, (B) 122-137 bp and (C) 162-188 bp for pl-iPSC and NPC. Plots show the average distribution of nucleosomes centred at and surrounding the RE1 site (n=871). Peaks correspond to the NRSF/REST complex, and nucleosomes in B and C respectively. D: Cluster analysis of frequency distributions centred on the NRSF/REST peak present from 122-137 bp size class of sequencing reads from pl-IPSC. Average sequencing read mid-point frequency distributions centred on the RE1 site ± 300 bp were plotted for the strongest (solid line) and weakest (dotted line) cluster. The same sites in each pl-IPSC cluster were used in corresponding plots for NPC. E: Schematic of nucleosome patterning at the RE1 site in pl-iPSC and NPC.

To separate the NRSF/REST footprint from that of the flanking nucleosomes, we constructed a nucleosome map using only larger fragment sizes (162-188bp), minimising the overlap with the smaller 122-137 bp fragments (Figure 3C). This shows a complete separation of NRSF/REST binding from nucleosome positions, and significantly that the average pattern of nucleosome positioning is unchanged in NPC in the absence of the NRSF/REST complex. To ensure that genome averaging did not mask any differences at these sites, we carried out cluster analysis (Figure 3D) to establish that 64% of the selected RE1 sites had detectable peaks, corresponding to NRSF/REST complex in undifferentiated pluripotent cells, but absent at the NPC stage. In pluripotent cells, there was a strong correlation between the presence of NRSF/REST at the RE1 binding site and an array of well-positioned nucleosomes. This nucleosome array pattern was retained at the same sites in NPC cells, despite the absence of the NRSF/REST complex. These observations indicate that NRSF/REST binding sites are associated with a distinct consensus nucleosome positioning pattern, however this pattern is not dependent on the presence of high levels of the NRSF/REST protein complex (Figure 3E). The function of this retained chromatin structure is not known. However, it is possible that this structure might persist so that the protein complex can rapidly re-engage in the neuronal state, where it is normally absent. It is known that REST levels can increase in later development or in response to cellular stress, and targets a subset of genes that are poised to respond to changes in REST levels [22,23].

Our analysis of chromatin states suggests a strong enrichment of nucleosomes at CTCF sites (chromatin state 8). To probe this interaction further, we derived 9,478 CTCF sites based on mapping the CTCF binding motif [24] with ChIP-seq data from human stem cells (H1ESC). 67% of these CTCF sites are present within chromatin state 8, representing a 600-fold enrichment (Figure 2A). Again there was a distinct peak in our small particle data (122-137 bp) from pluripotent iPSC (Figure 4A), corresponding to CTCF binding. This peak was significantly reduced in the NPC cells, but not completely absent. Cluster analysis demonstrated that this peak did not arise from averaging a smaller number of sites with strong peaks, but in fact represented a decreased occupancy at all CTCF sites during the transition from pluripotent cells to NPC (Figure 4B).

**Figure 4:**
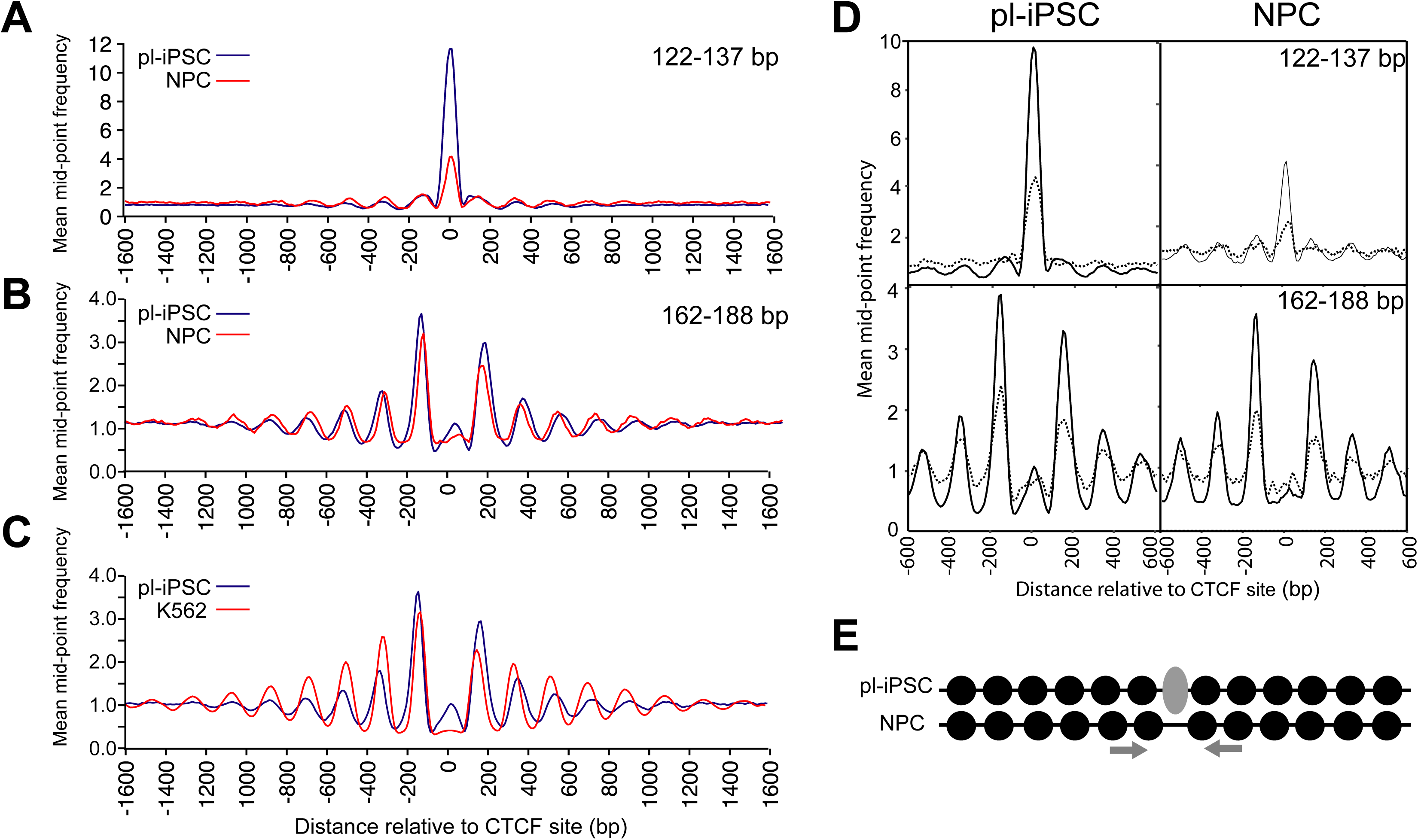
Nucleosome patterning associated with the CTCF binding sites. A: Average frequency distribution for sequence read mid-point data in the size range of 122-137 bp centred on the CTCF binding site. This shows a distinctive peak corresponding to the CTCF protein complex, which is present in pl-iPSC, but reduced in NPC. B: Average frequency distribution for sequence read mid-point data centred on the CTCF binding site in the size range of 162-188 bp, corresponding to larger nucleosome footprints, for pl-iPSC and NPC cells. This shows that the pattern of nucleosome positioning is independent of the amount CTCF protein complex. Although positioned nucleosomes are retained flanking CTCF sites, their positions are shifted closer to the CTCF site and their spacing is altered. C: Average frequency distribution for nucleosomes (sequence read mid-point data) centred on the CTCF site for pl-iPSC (162-188 bp) and K562 (total MNase-seq data) cells. Again positioned nucleosomes are retained flanking CTCF sites in K562 cells, but with shifted positions and altered spacing. D: Cluster analysis of frequency distributions centred on the CTCF peak present from 122-137 bp size class of sequencing reads from pl-IPSC. Average sequencing read mid-point frequency distributions centred on the CTCF site ± 300 bp were plotted for the strongest (solid line) and weakest (dotted line) cluster. The same sites in each pl-IPSC cluster were used in corresponding plots for NPC. E: Schematic of nucleosome patterning at the CTCF site in pl-iPSC and NPC.

Nucleosome positioning data revealed an array of positioned nucleosomes flanking the CTCF site in pluripotent hiPS, and again, in a similar fashion seen for NRSF/REST, these were retained in NPC cells, despite the decrease in CTCF binding (Figure 4 B&C). Furthermore, our analysis of the MNase-seq dataset for K562 cells showed a similar nucleosome patterning surrounding the CTCF sites of these lymphoblastoid cells (Figure 4D) [7]. However, close inspection of the nucleosome positioning pattern showed that the spacing of nucleosomes both upstream and downstream of the CTCF site in NPC was altered and that they were positioned nearer to the CTCF site than in pluripotent cells (Figure 4 B–D). This indicates that minor re-positioning of nucleosomes flanking CTCF sites occurs during the early stages of neural cell differentiation. A recent report using an alternative method based on Histone 3 ChIP-seq also showed a nucleosome array centred on CTCF of human myeloid leukaemia cells (HL60) and altered nucleosome spacing following retinoic acid induced differentiation [25]. In *Dictyostelium*, loss of ChdC, a type III CHD family chromatin remodeler, changes nucleosome spacing, indicating a capacity to specifically regulate spacing within nucleosomes [5]. In mammals, CTCF has been shown to complex with CHD8, an orthologue of ChdC [26], suggesting a mechanism for nucleosome translocation at these sites. Although apparently minor, small changes in nucleosome spacing are likely to have profound changes in the local chromatin structure [27] and given the role of CTCF in determination of the Topological Associated Domain (TAD) structures, this may lead to quite significant changes of high level chromatin architecture and hence gene regulation.

In conclusion, we have used genome wide mapping of human iPSC to follow the dynamics of nucleosome positioning from the pluripotent state to an early stages of neuronal development. In the pluripotent cell, only a small proportion of nucleosomes are highly-positioned, with the positions of more than 99% of nucleosomes showing considerable variation across the cell population. During differentiation to NPC, the number of highly positioned nucleosomes increases substantially to a number that approximates to that found in other differentiated cell lines. However, there are very few organised arrays of nucleosomes with the genomes of these human cell types, and no strong correlation of nucleosome positioning to active or repressive chromatin or gene activity. Remarkably, where nucleosome arrays do occur, they mark chromatin structures that are retained during cell differentiation, and may represent defined regulatory sites controlling chromatin accessibility, modifications or higher order 3D organisation [28]. We propose that these sites where positioned nucleosomes are retained during development form important regulatory nodes within the dynamic chromatin architecture – the 4D nucleome.

## SUPPLEMENTAL INFORMATION

Supplemental information contains one Figure and four Tables.

## ACKNOWLEDGMENTS

The research reported in the manuscript was supported by grant from The Waterloo Foundation to A.J.H and a BBSRC-funded Daphne Jackson fellowship awarded to J.C.H.

## AUTHOR CONTRIBUTIONS

J.C.H, N.D.A. and A.J.H. designed the experiments; J.C.H. and N.A.K. generated the data; J.C.H and A.J.H performed the analysis and wrote the manuscript.

## METHODS

### DATA AND SOFTWARE SHARING

Raw and processed sequencing read data is available on GEO: https://www.ncbi.nlm.nih.gov/geo/query/acc.cgi?acc=GSE117870

All the scripts used in this analysis are available on request.

#### Cell culture

The 34D6, male, human iPS cell line (a gift from Prof Chandran, Edinburgh) [29] was cultured in mTeSR1 (Stem Cell Technologies) following the manufacturer’s instructions. Tissue culture plates (Nunclon, Invitrogen) were coated for at least 2 hours with MatrigelTM (BD Biosciences,VWR) diluted 1:75 with DMEM/F12 (Invitrogen). 34D6 iPS cells were plated at a density of ∼10^6^ cells/10cm plate in complete mTeSR1TM medium containing 10µM Y27632 (Tocris) and incubated at 37°C in a standard 5% CO2, humidified incubator (Binder). To passage iPS cells, cells were first treated with 10µM Y27632 for 2hrs, then washed with Ca^2+^/Mg^2+^-free PBS and cell colonies lifted from the plate by incubation with Dispase (Stem cell technologies) containing 10µM Y27632 (Tocris). Colonies were fragmented by gentle trituration, collected by centrifugation (180 x g) and re-suspended in medium for re-plating.

#### Dual SMAD inhibition

For neural differentiation a dual SMAD inhibition protocol was used [30]. 34D6 iPS cells were harvested as described above for iPS cell passaging. 34D6 iPS cell colony fragments were plated in non-adherent bacteriological grade culture dishes in ADF differentiation medium to allow for embryoid body formation [31]. ADF differentiation medium comprised advanced DMEM/ F12TM medium (Invitrogen) supplemented with penicillin/streptomycin (5µg/l, Invitrogen), L-glutamine (200mM, Invitrogen), 1x Lipid concentrate (Invitrogen), 7.5µg/ml holo-transferrin (Sigma), 14µg/ml Insulin (Merck), 10µM β-mercaptoethanol (Sigma). Medium was supplemented with 10µM Y27632 (Tocris) for the first 2 days, with 10µM SB-431542 (Tocris) until day 4 of differentiation and 0.5µM LDN193189 (Miltenyi) until day 8 of differentiation. Medium was changed every 2 days. At day8, neuralised embryoid bodies were washed with Ca^2+^/Mg^2+^-free PBS and then dissociated by incubation at 37°C with accutase (PAA laboratories). A single cell suspension was obtained by gentle trituration and cells washed with ADF medium and harvested by centrifugation at 1000rpm. Neural progenitors were then plated onto tissue culture plates coated with 0.1µg/ml Poly-L-Lysine (Sigma) and 10µg/ml Laminin (Sigma) in ADF medium + 5ng/ml FGF2. 34D6-derived neural progenitors were grown to sub-confluency and passaged once by dissociation with accutase and re-plating. The 34D6 iPS cells derived neural progenitors were harvested for nucleosome preparation on day 16 of differentiation. The NPC cell population was validated using immunocytochemistry by checking for the presence of the NPC-specific markers, such as Nestin and the loss of pluripotency markers, for example Oct-4. Greater than 90% of the differentiated population contained neural progenitors.

#### *In vivo* MNase digestion of chromatin

Chromatin was prepared from three bio-replicates of human iPS cells and the same cells differentiated to NPC cells as described previously for the *S. cerevisiae* genome [4]. The cell membranes and nuclei were made permeable to MNase using NP-40 [32,33] and each bio-replicate was treated by *in vivo* digestion with 300U/ml MNase at room temperature for 4 minutes. DNA fragments were purified from each MNase treated sample, then the samples for each cell type were pooled in equimolar amounts. DNA extracted from chromatin samples was size-fractionated on agarose gels. 25-35ug of DNA less than 300 bp was size-selected for each cell type.

#### Paired end mode DNA sequencing

All of the DNA fragments less than 300 bp in size were utilised for paired-end mode sequencing by Source BioScience (http://www.sourcebioscience.com/) on an Illumina HiSeq 2000 platform (HiSeq) using a read-length of 50 bp. Eight flow cells were used for each cell type to obtain a sufficient depth of coverage of the human genome. A standard Illumina paired-end mode sequencing protocol was used, apart from the omission of the nebulisation step and the addition of a further gel purification step to eliminate any excess concatenated linkers after the ligation of linker DNA to the sample. Base calling and quality control of the sequencing data was performed using Real Time Analysis (RTA) 1.09, CASAVA 1.8 software.

#### Alignment of paired-end reads to the genome

A total of 3.4 and 3.0 billion paired-end reads were obtained in fastq format from iPS and NPC cells respectively and aligned to the human genome assembly hg19 using Bowtie version 0.12.8 [34] The command line options for bowtie were as follows: bowtie -v 3 --trim3 14 --maxins 5000 --fr -k 1 --best -p 12.

#### Creation of nucleosome maps

Subsequent data processing to create nucleosome maps from iPS and NPC cells was undertaken in a similar manner to the method previously described for the *S. cerevisiae* genome [4]

#### Data validation and normalisation

The number of reads obtained for each human autosome in each cell type was determined and utilised for further analysis. To compensate for the slight genome-wide difference in total read counts between the two cell types, the read counts for the NPC autosomal genome were multiplied by the ratio of the total aligned reads in the autosomal genomes of iPS v NPC cells, which was 1.117.

#### Read mid-point frequency distributions

Paired-end reads obtained from Bowtie alignments in .sam format were sorted into separate chromosome-specific files. To represent a unique position for each paired-read, the genomic position of the mid-point of the insert DNA was calculated, representing a position equivalent to the nucleosomal dyad [35]. The reads were separated and filtered into three size classes:112-137 bp, 138-161 bp (nucleosomes), 161-188 bp. The frequency distribution for the mid-point position of the sequencing reads was derived at 10 bp resolution and the data was smoothed using a 3bin moving average. Frequency distributions were output as chromosome-specific files in .sgr format: chromosome identification: chromosomal location of the start of each 10 bp bin: frequency of the paired-read midpoint values that fall within that 10 bp bin.

#### Published human nucleosome maps

Published human nucleosome maps [7] from the K562 chronic myelogenous leukaemia (Cml) cell line [36] and from the B-lymphoblastoid cell line GM12878 (Corriell biorepository) were converted from bigwig to bedgraph format and the chromosome-specific bedgraph files were then converted to .sgr files by binning the data into 10 bp bins and calculating a 3 bin moving average exactly as described for the iPS and NPC data above. The final re-processed maps were validated as for the iPS and NPC maps, generating the total numbers of aligned paired - end reads for each genome.

#### Locating patterns of positioned nucleosomes

In order to locate and quantify the number of highly positioned nucleosomes, a heuristic peak-finding algorithm was developed. Peaks were defined as three consecutive 10 bp bins where the value of the paired-read midpoint frequency in the central 10 bp bin was between 30 and less than 1000. The lower threshold for the paired-read midpoint frequency values for the bins either side of the central bin was set at two (the ‘noise’ threshold). The upper threshold for each of the paired-read midpoint frequency values for the bins either side of the central bin, was less than the paired-read midpoint frequency in the central bin. The upper threshold was chosen to exclude regions of the genome with high frequencies of paired-read midpoint values found at the ends of chromosomes and at runs of repeats and at centromeres.

Peaks in the genomic distribution of sequence read mid-points, were given explicit genome positions using Peakfinder for all of the nucleosome maps: iPS, NPC and K562.

#### Detecting nucleosome arrays

An in-house python script was used to calculate the distance between the genomic locations of all highly positioned nucleosomes across the genome for each cell type, iPS, NPC and K562. Positioned nucleosomes 150-200bp apart were located and those existing as singletons or in arrays of 2,3 or more nucleosomes were counted.

#### Nucleosome positions in different cell types

In house perl scripts were used to compare a) the genome-wide overlap in the locations of positioned nucleosomes within ± 10 bp in iPS and NPC cells (Fig 1B).

#### Chromatin states

Chromatin state data file: wgEncodeBroadHmmH1hescHMM.bed containing chromatin state information derived from H1ESC cells was downloaded from http://genome.ucsc.edu/cgi-bin/hgFileUi?db=hg19&g=wgEncodeBroadHmm. For iPS cells we aligned the genomic positions of the peaks in the nucleosome map determined using the heuristic PeakFinder with the chromatin state regions determined in H1ESC cells. Thus we calculated genome-wide total for the number of nucleosomes in the following chromatin states [8,9] : Active promotor = state 1; Insulator = state 8; Repressed = state 12 Heterochromatin = state 13. We calculated the expected number of nucleosomes in any particular chromatin state, assuming that positioned nucleosomes are dispersed randomly across the genome (Tables S2). Similarly, we determined the chromatin state for the CTCF sites we derived (using the H1ESC ChIP-seq data combined with the CTCF binding motif) and calculated the expected chromatin states for CTCF sites assuming that their distribution across the genome is random (Tables S3).

#### Construction of genomic feature lists

The positions of transcriptional start sites for all of the full-length transcripts for human genome build Feb 2009 GRCH37/hg19 were derived as follows. Track = Gencode Genes v17, table = basic, was downloaded from the UCSC genome browser (contains 94,151 TSS). From this a list of non-redundant, strand-specific autosomal TSS was derived (n= 83,179) using in-house scripts.

Expression data from ESC cells and ESC cells differentiated to the N2 stage of development [12] was used to generate two lists of genes expressed specifically in a) ESC cells (n = 3833) and b) N2 cells (n =2082). The minimum expression threshold was 0.1 (normalised RPKM) and the difference in expression was at least two-fold difference between ESC and NPC in each case.

TSS of genes expressed specifically in a) ESC cells and b) N2 cells were tested for the presence of positioned nucleosomes within ± 300bp of the TSS by mapping the genomic locations of highly positioned nucleosomes generated by the PeakFinder tool to a window ± 300bp of the TSS in each case.

#### Transcription factor binding motifs

The genomic positions of the consensus binding sequence for each transcription factor were extracted from fasta files from the Feb. 2009 assembly of the human genome (hg19, GRCh37 Genome Reference Consortium Human Reference 37 (GCA_000001405.1)) using in-house perl scripts. Fasta files were downloaded from http://hgdownload.cse.ucsc.edu/goldenPath/hg19/chromosomes/

The consensus binding motif used in this study for each transcription factor is shown in Table S4. The consensus binding motif for YY1, M1, from factorbook derived from ChIP-seq data derived from H1ESC cells was used. (http://www.factorbook.org) [37] The ATF2 CRE binding motif was taken from Hai et al [38]. The PAX6 consensus binding sequence was derived using (ChIP) in ES-derived neuroectodermal cells (NECs) [39]. CTCF binding positions were derived by mapping the consensus binding motif derived by Ong *et al* [24] with H1ESC ChIP region data from the Broad institute downloaded from the UCSC genome browser, file wgEncodeAwgTfbsBroadH1hescCtcfUniPk.narrowPeak (UCSC accession wgEncode EH000085). RE1 sites were derived by mapping the consensus binding motif derived by Bruce *et al* [18] with ChiP data that was derived from the ENCODE database [19]. ChiP data was downloaded using the UCSC genome browser from human genome assembly Feb 2009 GRCH37/hg19 Group=regulation, Track name = TXnFactorChiP, Table= wg EncodeRegTfbsCLusteredv2 from uniform processing of data from the Jan. 2011 ENCODE data freeze.

#### SiteWriter

The in-house perl script SiteWriter [4] was used to construct average frequency distributions of the sequencing read mid-point values at and surrounding genomic feature loci within a user-defined window, for example at transcription factor binding sites, and surrounding positioned nucleosomes (Fig 1C). The output from the SiteWriter script comprises two files: a) a CFD.txt file of the normalised average frequency values. The values are normalised by dividing the frequency value in each bin by the number of bins specified in the user-defined window. b) a C3.txt file which contains a matrix of locally-normalized dyad frequency values for every bin position. The C3.txt file data were used in cluster analysis.

#### Cluster analysis

Cluster analysis was undertaken in R, using the Canberra method to generate a distance matrix from the C3.txt file from the output of the SiteWriter script. Dendrograms generated by hierarchical agglomerative clustering were used to determine the number of groups to use in k-means clustering. Cluster data for genomic features were used to construct average frequency distributions of the read mid-point values at and in the bins surrounding genomic feature loci within a user-defined window for selected clusters.

